# Interleukin-36 promotes systemic Type-I IFN responses in severe psoriasis

**DOI:** 10.1101/496851

**Authors:** Marika Catapano, Marta Vergnano, Marco Romano, Satveer K Mahil, Siew-Eng Choon, A David Burden, Helen S Young, Ian M Carr, Helen J Lachmann, Giovanna Lombardi, Catherine H Smith, Francesca D. Ciccarelli, Jonathan N Barker, Francesca Capon

**Author notes:** These authors contributed equally (Marika Catapano, Marta Vergnano).

## Abstract

Psoriasis is an immune-mediated skin disorder associated with severe systemic co-morbidities. Both chronic and acute forms of the disease are characterised by abnormal interleukin (IL)-36 signalling. While the mechanisms whereby IL-36 promotes cutaneous inflammation are well established, its systemic effects have not been investigated. To address this issue, we initially measured leukocyte gene expression in generalised pustular psoriasis, an acute disease variant caused by mutations of the IL-36 receptor antagonist. By undertaking whole-blood and neutrophil RNA-sequencing in affected individuals, we identified a Type-I IFN signature, which correlated with IL-36 signalling up-regulation. We then validated these observations in patients with chronic plaque psoriasis. Finally, we demonstrated that IL-36 acts directly on plasmacytoid dendritic cells, where it potentiates Toll-like Receptor (TLR)-9 activation and IFNα production. This effect was mediated by the induction of PLSCR1, an endosomal TLR-9 transporter. These results define an IL-36/TLR-9/Type-I IFN axis that could be targeted for the treatment of psoriasis co-morbidities.

## INTRODUCTION

Interleukin-36α, -β and-γ (hence IL-36) are group of IL-1 family cytokines that are mainly produced by keratinocytes, monocytes and dendritic cells^1^. IL-36 signalling plays an important role in epithelial immune homeostasis and its de-regulation has been repeatedly implicated in the pathogenesis of plaque psoriasis (PsV), a common and chronic, immune-mediated skin disorder^1^.

Numerous studies have shown that IL-36 responses are elevated in PsV skin^2, 3, 4, 5^. Moreover, we have demonstrated that IL-36 stimulates chemokine production and amplifies the effects of IL-17 signalling in psoriatic lesions^3^. Finally, animal studies have established that IL-36 promotes the activation of dendritic cells and the polarization of T lymphocytes into Th17 cells^6^. Thus, the mechanisms whereby IL-36 contributes to cutaneous inflammation have been extensively investigated. Its systemic effects, however, remain poorly understood.

We and others have shown that recessive mutations of the IL-36 receptor antagonist (*IL36RN*) are associated with generalised pustular psoriasis (GPP), a disease variant characterized by severe extra-cutaneous symptoms^7, 8^. In fact, GPP patients suffer from flares of skin pustulation that are often accompanied by acute systemic upset (fever, elevation of acute phase reactants and neutrophilia)^9^. This suggests that IL-36 signalling is likely to influence immune responses beyond skin.

Of note, extra-cutaneous co-morbidities are also well documented in PsV, as individuals suffering from severe disease are at high risk of psoriatic arthritis, metabolic syndrome and atherosclerosis^9, 10, 11^. It has therefore been proposed that PsV is a systemic disease, manifesting with skin, joint and vascular inflammation^12, 13^.

In this context, we hypothesise that abnormal IL-36 signalling has extra-cutaneous effects in both GPP and PsV, driving acute systemic flares in the former and contributing to a state of chronic systemic inflammation in the latter. To explore this model, we have integrated the transcription profiling of patient leukocytes with the characterisation of IL-36 responses in circulating immune cells. Our experiments show that IL-36 potentiates Toll-like receptor (TLR)-9 activation and enhances Type-I IFN production by plasmacytoid dendritic cells (pDCs). Thus, we have identified an IL-36/TLR-9 axis which up-regulates Type-I IFN, a cytokine that has been repeatedly implicated in the pathogenesis of systemic immunity, arthritis and atherosclerosis.

## RESULTS

### Expression profiling identifies a Type-I IFN signature in GPP and PsV whole-blood samples

We reasoned that GPP would represent an ideal model in which to investigate the systemic effects of IL-36, given the well-established link with *IL36RN* mutations^7, 8^ and enhanced IL-36 activity^14^. We therefore undertook RNA-sequencing of whole-blood samples obtained from 9 affected individuals and 7 healthy controls (Supplementary Table 1). This identified 117 differentially expressed genes (DEG); 111 of which were up-regulated (fold change≥1.5) at a False Discovery Rate (FDR) < 0.05 (Fig. 1a and Supplementary Data 1). As expected, key IL-36 target genes (*IL1B, PI3, VNN2, TNFAIP6, SERPINB1*)^15^ collectively showed a significant up-regulation in cases vs. controls (*P*=0.019) (Fig.1b). Of note, the analysis of a publicly available PsV dataset (whole-blood samples obtained from 33 cases vs 44 controls)^16^ identified a moderate, but statistically significant, over-expression of the same genes (*P*=0.001) (Fig.1b), providing the first indication that IL-36 may have systemic effects in PsV.

**Figure 1.**
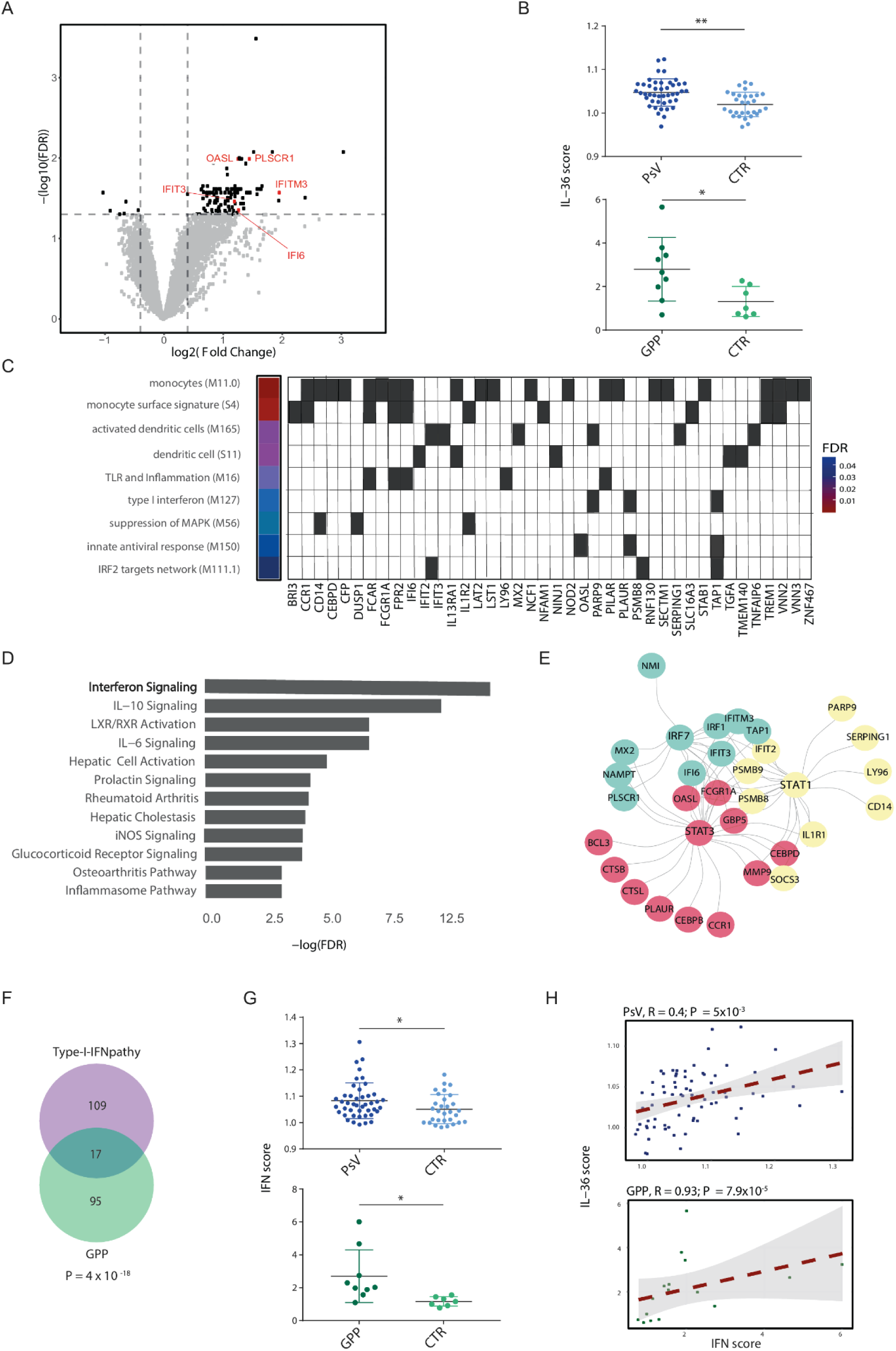
Transcription profiling of GPP and PsV whole-blood samples uncovers a Type-I IFN signature that correlates with the over-expression of IL-36 dependent genes. (**A**) Volcano plot showing the genes that are differentially expressed in GPP patients (black dots). The horizontal and vertical dashed lines represent significance (FDR <0.05) and fold change (FC>1.5; FC<0.7) thresholds, respectively. The genes used to measure the IFN score are labelled in red font. (**B**) Dot plots showing higher expression of IL-36 dependent genes in whole-blood samples of GPP (green) and PsV (blue) patients, compared to healthy controls (CTR). The IL-36 score for each individual was defined as the median expression of the *IL1B, PI3, VNN2, TNFAIP6,* and *SERPINB1* genes. The data for each patient and control group are presented as mean +/-standard deviation; \**P*<0.05, \*\**P*<0.01 (unpaired t-test). (**C**) Diagram showing the most significantly enriched transcriptional modules among the genes that are up-regulated in GPP. The heat map on the left reports the FDR associated with each module, with the underlying up-regulated genes shown on the grid as grey cells. (**D**) Bar plot illustrating enriched pathways (FDR<0.05) detected among the genes that are over-expressed in GPP (Ingenuity Pathway Analysis). (**E**) Upstream regulatory network showing that IRF7, STAT1 and STAT3 drive the up-regulation of numerous genes that are over-expressed in GPP (Ingenuity Pathway Analysis). (**F**) Venn diagram showing a significant overlap between the genes that are up-regulated in GPP (green) and Type-I-IFN driven autoinflammatory disease (purple). The p-value was derived with a hypergeometric test. (**G**) Dot plots showing an elevated IFN score in whole-blood samples of GPP (green) and PsV (blue) patients, compared to healthy controls (CTR). The data are presented as mean +/- standard deviation; \**P*<0.05, \*\**P*<0.01 (unpaired t-test). (**H**) Scatter plots showing that IL-36 and IFN scores are significantly correlated, in both GPP and PsV patients. Dashed regression lines are plotted with their 95% confidence intervals (grey areas).

To further explore the biological significance of our findings, we mapped the genes up-regulated in GPP to the co-expression modules described by Li et al^17^, which define well characterised features of immune function. We found that the over-expressed genes were significantly enriched among modules related to innate immune activation (e.g. *enriched in activated dendritic cells*, FDR<0.005) and antiviral responses (e.g. *viral sensing*, and *type I IFN response*; FDR<0.05 for both) (Fig. 1c). These findings were validated by Ingenuity Pathway Analysis (IPA), which identified *interferon signalling* as the most significantly enriched pathway (FDR<5×10^−6^) (Fig. 1d and Supplementary Data 2). An upstream regulator analysis also highlighted IRF7, STAT1 and STAT3 as the transcriptional activators that are most strongly associated with gene over-expression (FDR<10^−10^ for all) (Fig. 1e and Supplementary Data 2). This is of interest since STAT1 and STAT3 are critical mediators of IFN signal transduction, and IRF7 is a key driver of IFN-α production in pDCs^18^.

Finally, the analysis of a publicly available dataset^19^ demonstrated a significant overlap (*P*=4×10^−18^) between the genes that are up-regulated in GPP and those that are over-expressed in autoinflammatory syndromes caused by excessive Type-I IFN production (Fig. 1f). Of note, no such overlap was found with the up-regulated genes detected in cryopyrin associated periodic syndrome (CAPS) ^20^, a disease caused by excessive IL-1 activity, which was analysed as a negative control (Supplementary Fig. 1a). Thus, the presence of a Type-I IFN signature in GPP leukocytes is supported by several lines of evidence.

To further investigate the relevance of these observations we built an interferon score (IS) by measuring the aggregate expression of 5 Type-I IFN regulated genes (*IFI6, IFIT3, IFITM3, OASL, PLSCR1*). As expected, the score was elevated in GPP cases, compared to controls. A similar increase was observed in the publicly available PsV dataset (Fig. 1g), indicating that abnormal Type-I IFN activity is also present in PsV leukocytes. Importantly, we found that the IS documented in GPP and PsV significantly correlated with the IL-36 signatures observed in these datasets (*P<*0.01) (Fig. 1h). Thus, we have shown that psoriatic patients display abnormal Type-I IFN activity at the systemic level, which may be linked to increased IL-36 production.

### The Type-I IFN signature is readily detectable in GPP neutrophils

The presence of heterogeneous cell populations in whole-blood samples can complicate the interpretation of transcription profiling experiments. We therefore sought to validate our results through an independent analysis of a single cell type. Given that neutrophils play a critical role in systemic inflammation and can be activated by Type-I IFN signalling^21, 22, 23^, we focused our attention on these cells. We obtained fresh blood samples from 8 GPP cases and 11 controls. We then isolated highly pure (>95%) neutrophil populations (Supplementary Fig. 2a and Supplementary Table 1). The RNA-sequencing of these samples detected all the genes (n=3,133) that are transcribed in the neutrophils sequenced by the BluePrint consortium^24^, with very low expression levels observed for a further 252 genes originating from other cell types (<0.3 transcript per million (TPM) on average). This further validates the high purity of our neutrophil samples (Supplementary Fig. 2b).

**Figure 2.**
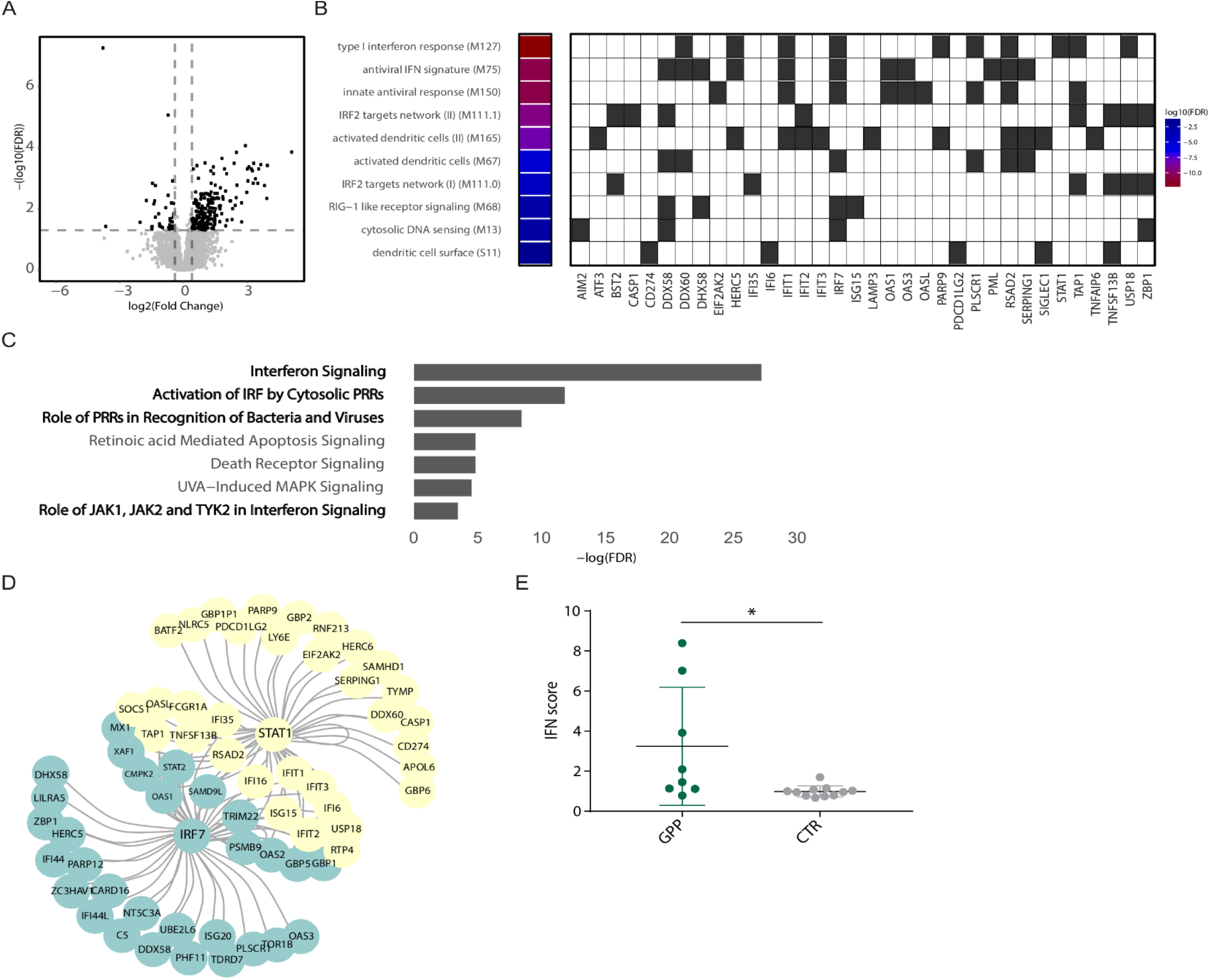
Transcription profiling of neutrophil GPP samples confirms the presence of a Type-I IFN signature. (**A**) Volcano plot showing the genes that are differentially expressed in GPP patients (black dots). The horizontal and vertical dashed lines represent significance (FDR <0.05) and fold change (FC>1.5; FC<0.7) thresholds, respectively. (**B**) Diagram showing the most significantly enriched transcriptional modules among the genes that are up-regulated in GPP. The heat map on the left reports the log10(FDR) associated with each module, with the underlying up-regulated genes shown on the grid as grey cells. (**C**) Bar plot illustrating the most significantly enriched pathways detected among the genes that are up-regulated in GPP (Ingenuity Pathway Analysis). IFN-related pathways are highlighted in bold font (**D**) Upstream regulatory network showing that IRF7 and STAT1 drive the up-regulation of numerous genes that are differentially expressed in GPP (Ingenuity Pathway Analysis). (**E**) Dot plots showing an elevated IFN score in the neutrophils of GPP patients, compared to healthy controls. The data are presented as mean +/- standard deviation; \**P*<0.05 (unpaired t-test).

Differential expression analysis detected 200 up-regulated (fold change≥1.5) and 31 down-regulated (fold change≤0.70) genes, at an FDR<0.05 (Fig. 2a and Supplementary Data 3). The analysis of transcriptional networks identified *Type-I interferon response* as the most significantly enriched module (FDR<10^−12^), followed by *innate antiviral response* and *antiviral interferon signature* (FDR<10^−10^ for both) (Fig. 2b). IPA also demonstrated a marked enrichment of pathways related to interferon signalling (FDR<10^−11^) (Fig. 2c and Supplementary Data 4) and highlighted IRF7 and STAT1 as the most likely drivers of gene up-regulation (FDR<10^-30^ for both) (Fig. 2d and Supplementary Data 4). In keeping with these findings, we found that GPP patients had an elevated IS compared to controls (*P*=0.02) (Fig. 2e). Thus, the analysis of purified neutrophils independently validated the Type-I IFN signature observed in whole-blood samples.

### The Type-I IFN signature can be validated in extended PsV and GPP datasets

We next sought to validate the type I IFN signature through the analysis of further affected individuals. We obtained whole-blood samples from 35 additional GPP patients and 7 healthy controls. We then measured the IS by real-time PCR of the *IFI6, IFIT3, IFITM3, OASL* and *PLSCR1* transcripts. The analysis of the expanded dataset (including the validation cohort as well as the nine samples that had been originally RNA-sequenced), confirmed the up-regulation of IFN stimulated genes (*P*=0.02) (Fig. 3a). We next examined neutrophils obtained from 17 GPP cases (including 7 from the RNA-seq dataset and 10 newly ascertained cases) and 16 PsV patients. To maximise the likelihood of observing a systemic Type-I IFN signature, we selected the PsV cases among patients suffering from severe disease (average Psoriasis Area and Severity Index (PASI): 17.9). We, however, excluded individuals who were being treated with TNF inhibitors, in order to avoid the confounding effects of TNF-dependent type-I IFN modulation^25^. Finally, we included three control groups in the analysis: 9 individuals affected CAPS, 13 cases of acral pustular psoriasis (a localised variant of the disease which manifests without systemic involvement) and 26 healthy volunteers. Real-time PCR demonstrated that the IS was significantly increased in GPP and PsV cases compared to healthy controls (*P*<0.005 for both). Conversely, and in keeping with the specificity of our observations, the IS of CAPS and acral pustular psoriasis patients were within the normal range defined in unaffected individuals (Fig. 3b).

**Figure 3.**
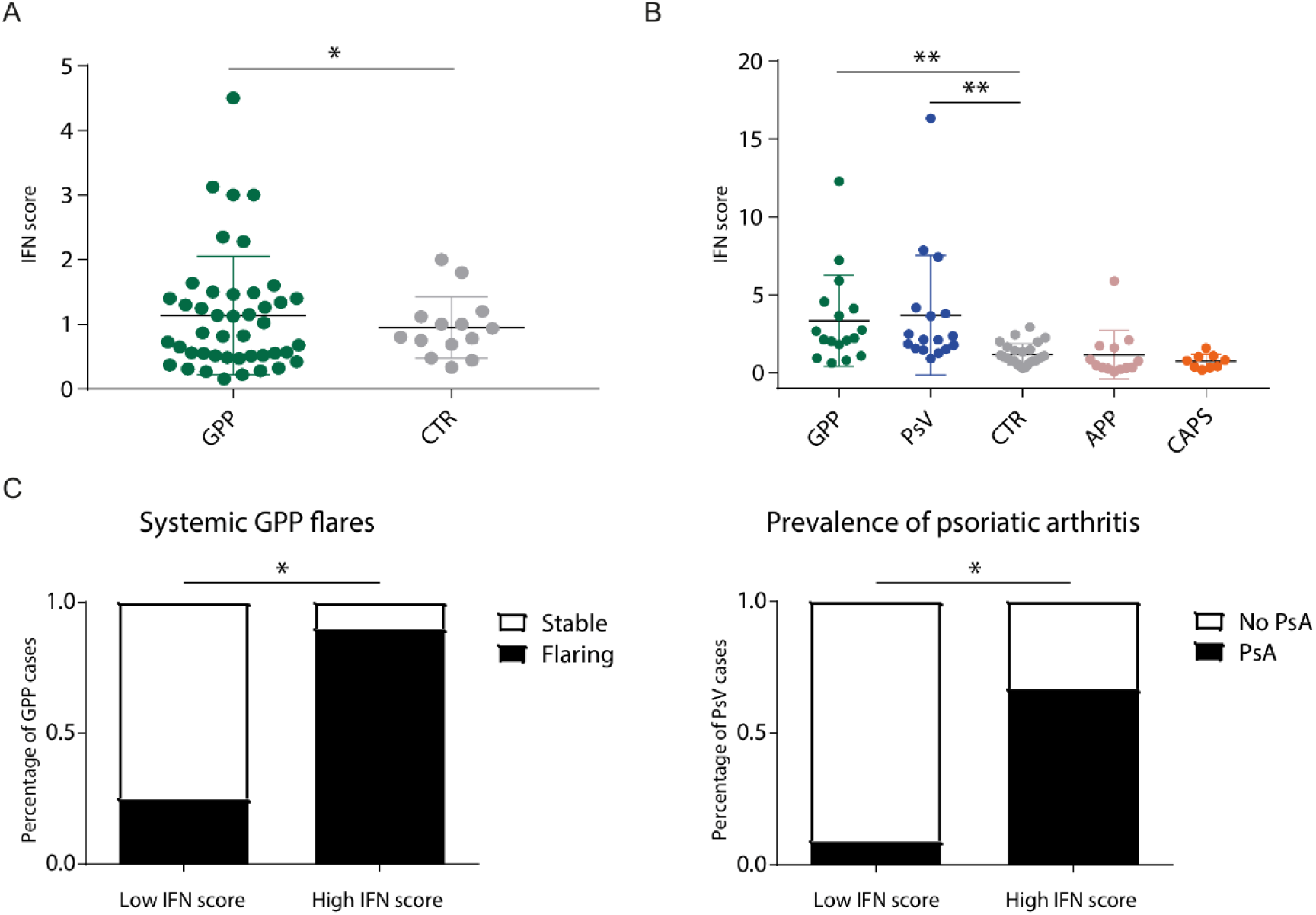
Validation of the Type-I IFN signature in extended datasets. (**A**) Dot plot showing an elevated IFN score in whole-blood samples of GPP patients (expanded dataset, n=44) compared to healthy controls (n=16). The data are presented as mean +/- standard deviation; \**P*<0.05 (unpaired t-test). (**B**) Dot plot showing an elevated IFN score in the neutrophils of GPP and PsV patients, compared to healthy individuals. Acral pustular psoriasis (APP) and CAPS cases were analysed as negative controls. The data are presented as mean +/- standard deviation; \*\**P*<0.01 (one-way ANOVA followed by Dunnett’s post-test). (**C**) Left panel: systemic flares are more prevalent in GPP patients with high interferon scores (n=8) compared to those with low interferon scores (n=9). Right panel: psoriatic arthritis (PsA) is more prevalent in PsV patients with high interferon scores (n=6) compared to those with low interferon scores (n=11). In both groups, the cut-off between high and low scores was defined as the median +2SD of the values observed in healthy controls. \**P*<0.05 (Fisher’s exact test).

Of note, a closer inspection of medical records showed that GPP patients with high IS were more likely to experience systemic flares than those with low IS (88% vs 33%; *P*=0.049). Likewise, high-IS PsV subjects more frequently suffered from psoriatic arthritis (80% vs 17%; *P*=0.03) (Fig. 3c).

Thus, the Type-I IFN signature detected by RNA-sequencing can be validated in independent PsV and GPP samples, where it is associated with features of systemic involvement.

### The IL-36 receptor is expressed on the surface of plasmacytoid dendritic cells

Having observed a marked up-regulation of IFN signature genes in both GPP and PsV, we hypothesised that IL-36 has a direct effect on Type-I IFN producing cells. To investigate this possibility, we systematically examined the surface expression of the IL-36 receptor (IL36R) in innate immune cells. We obtained peripheral blood mononuclear cells (PBMCs) and neutrophils from healthy donors and GPP patients. We then measured IL36R levels by flow-cytometry (Supplementary Figure 3). In keeping with published findings^26^, we found that IL36R was barely detectable on the surface of healthy neutrophils (Fig. 4a). We also showed that receptor levels were low in innate lymphoid cells (Fig. 4b), in classical, intermediate and pro-inflammatory monocytes (Fig. 4c), and only marginally higher in myeloid dendritic cells (mDC) (Fig 4d). Finally, we observed robust receptor expression in pDCs (Fig. 4d), a phenomenon that was especially noticeable among the GPP samples (Fig. 4e). Thus, we have shown that IL36R is preferentially expressed in pDCs, the cell type that produces the largest amounts of Type-I IFN.

**Figure 4.**
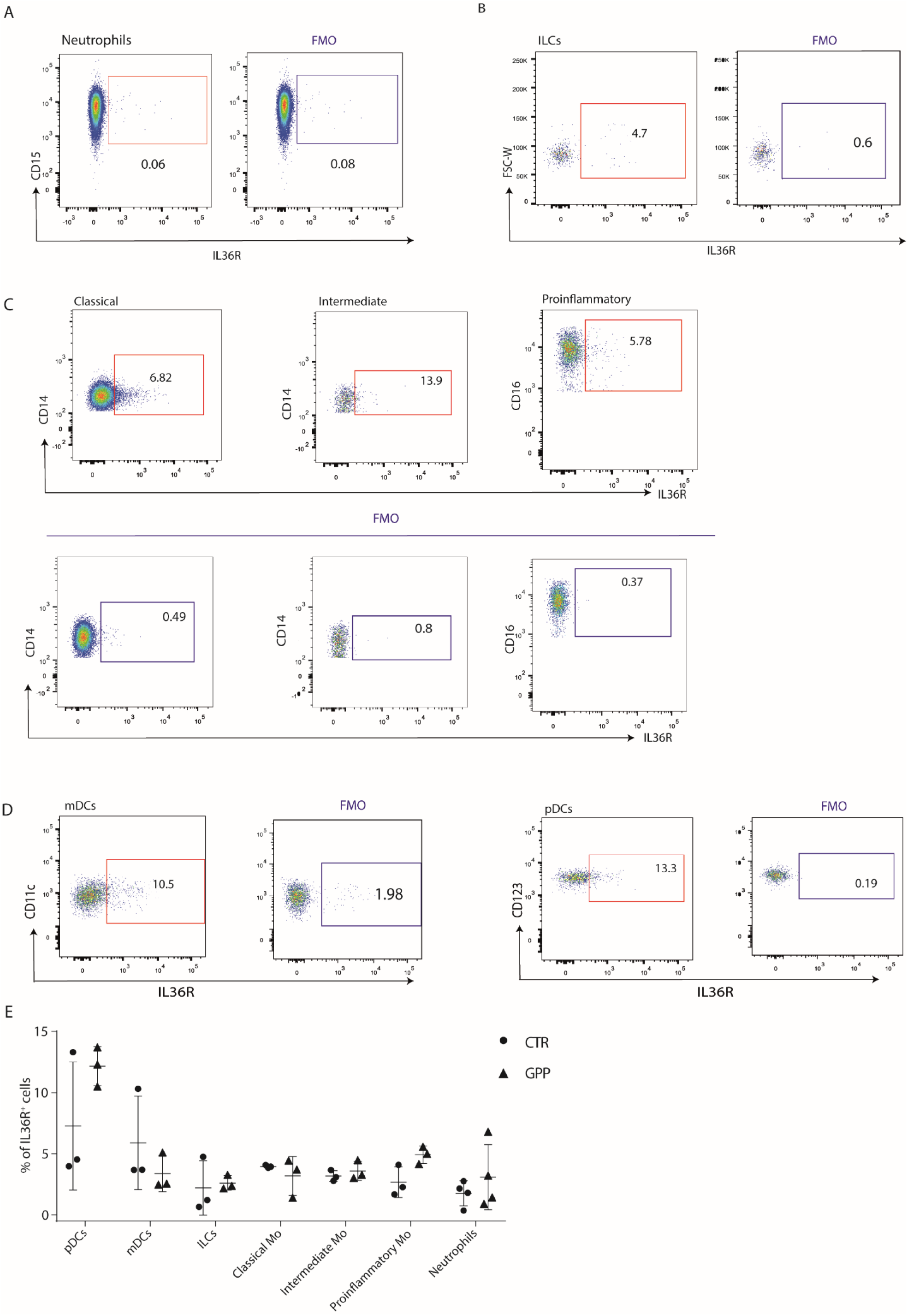
The IL-36 receptor is preferentially expressed by pDCs. (**A-E**) Representative flow cytometry analyses of IL36R surface expression in innate immune cells. Fluorescence minus one (FMO) controls are shown for each population. The following cell types were analysed: (**A**) neutrophils (gated as CD14^−^, CD15^+^, CD16^+^ cells); (**B**) innate lymphoid cells (lineage^−^ (CD3^−^, CD4^−^, CD19^−^, CD20^−^, CD56^−^), CD127^+^); (**C**) monocytes (CD3^−^, CD20^−^, CD19^−^, CD56^−^) separated into classical (CD16^−^, CD14^high^), intermediate (CD16^+^, CD14^+^) and pro-inflammatory (CD16^high^, CD14^−^) populations; (**D**) pDCs (lineage^−^, HLADR^+^, CD123^+^, CD11c^−^) and mDCs (lineage^−^, HLADR^+^, CD123^−^, CD11c^+^). (**E**) Dot plot showing the percentage IL36R^+^ cells in each leukocyte population (n= 3 GPP cases and 3 healthy controls for PBMCS; n=4 GPP cases and 4 healthy controls for neutrophils). Data are presented as mean +/- SEM. No significant differences were observed between GPP cases and healthy donors.

### IL-36 potentiates IFN-α production in response to Toll-like receptor 9 stimulation

Based on the results obtained in the above experiments, we hypothesised that IL-36 potentiates Type-I IFN production by pDCs. To investigate this possibility, we pre-treated PBMCs obtained from healthy donors (n=3) with IL-36 or vehicle. We then stimulated the cells with CpG-containing DNA (hence CpG), a Toll-like receptor (TLR)-9 ligand which induces IFN-α release by pDCs. Finally, we measured the up-regulation of the IS genes as a readout of Type-I IFN production. As expected, CpG treatment increased the expression of most IS genes. Of note, this effect was more pronounced in cells that had been pre-incubated with IL-36 (*P*<0.05 for *IFIT3, OASL* and *PLSCR1*) (Fig. 5a). This observation was validated by direct ELISA measurements of IFN-α production, demonstrating that IL-36 pre-treatment enhanced the response to CpG (*P*<0.01) (Fig. 5b). Finally, flow cytometry showed an increased proportion of IFNα**^+^** pDCs among the cells that had been stimulated with IL-36 and CpG, compared to those that had been exposed to CpG alone (Fig. 5c). Thus, multiple experimental readouts support the notion that IL-36 up-regulates TLR-9 dependent IFN-α release.

**Figure 5.**
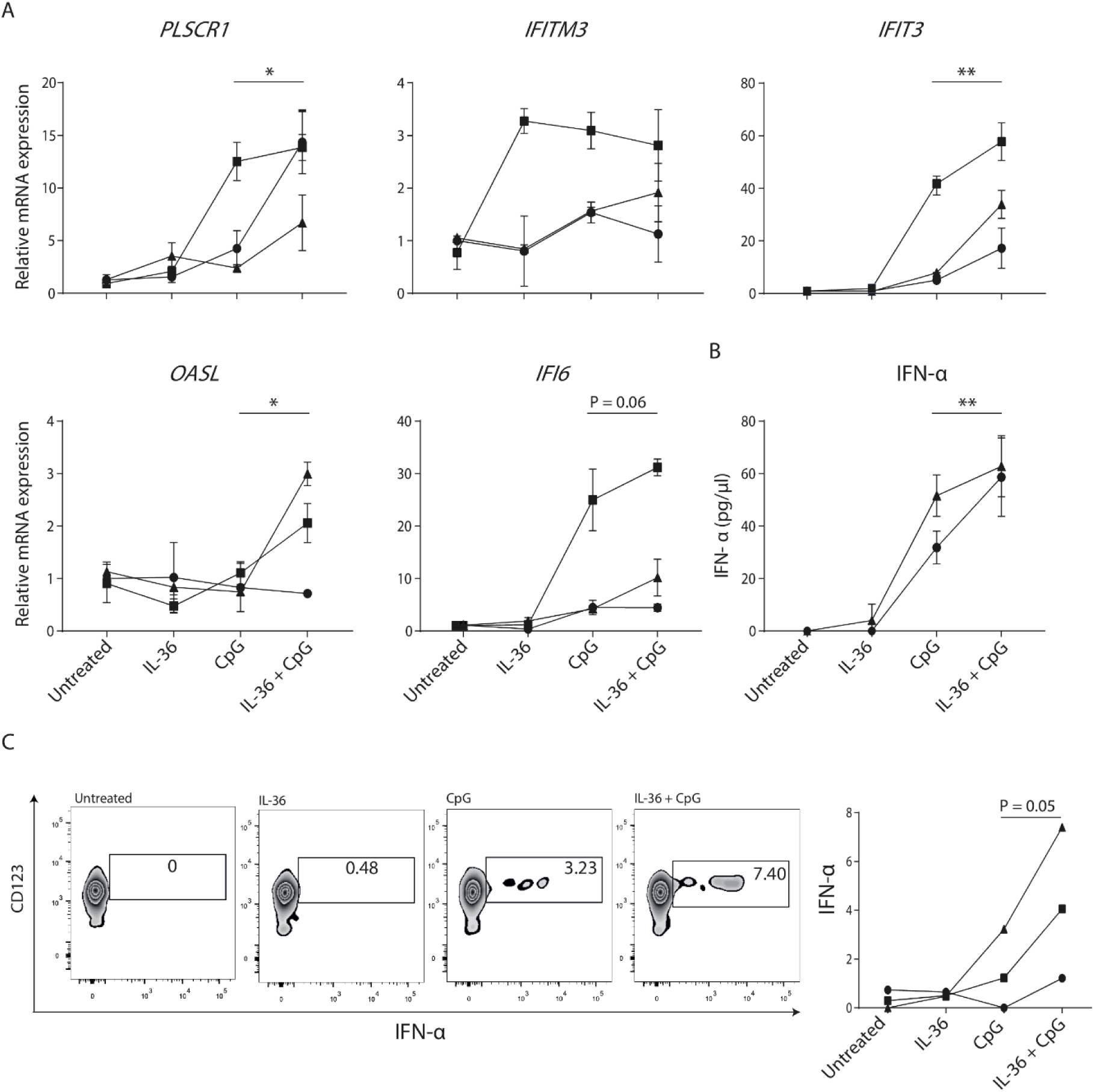
IL-36 enhances the production of IFN-α downstream of Toll-like receptor 9. Pre-treatment of PBMCs with IL-36 up-regulates CpG-dependent Type-I IFN production (**A**) PBMCs were stimulated with CpG for 6h, in the presence or absence of IL-36 pre-treatment (6h). The expression of the five interferon signature genes was then measured by real-time PCR and normalised to that of the *B2M* gene. Each line in the plot represents an independent healthy donor. Data points within a line represent mean +/- SD of results obtained in triplicate stimulations. (**B**) Following PBMC stimulation, IFN-α production was measured by ELISA. Each line in the plot represents an independent healthy donor. Data points within a line represent mean +/- SD of results obtained in triplicate stimulations. (**C**) Following PBMC stimulation, the percentage of IFNα^+^ pDCs was determined by flow cytometry. A representative set of zebra plots is shown on the left, while the panel on the right illustrates the data obtained in the individual healthy donors (n=3, each is represented by a line). \**P*<0.05; \*\**P*<0.01 (Friedman’s test, with Dunn’s post-test).

### IL-36 up-regulates *PLSCR1,* a known TLR-9 transporter

We next sought to define the mechanisms whereby IL-36 enhances IFN-α production downstream of TLR-9. A closer inspection of the PBMC stimulation results showed that IL-36 treatment can up-regulate the expression of *PLSCR1*, even in the absence of CpG. This is of interest, as the gene encodes phospholipid scramblase 1, a protein which has been shown to interact with TLR-9 and regulate its trafficking to the endosomal compartment^27^.

To further explore the link between IL-36 and PLSCR1, we first validated our initial observation through the analysis of additional donors (Fig. 6a). Next, we demonstrated that IL-36 treatment increases PLSCR1 protein levels in isolated pDCs, showing a direct effect of the cytokine on these cells (*P*<0.05) (Fig. 6b). Finally, we investigated the mechanism whereby IL-36 up-regulates *PLSCR1*. As expected for an IFN signature gene, an analysis of the *PLSCR1* promoter uncovered a STAT1 binding site. Given that IL-36 can signal through mitogen-activated protein kinases (MAPK)^1^, and that there have been reports of cross-talk between STAT1 and MAPK signalling^28^, we reasoned that the latter pathway was likely to be involved. Real-time PCR experiments confirmed this hypothesis, as the SB-203580 MAPK inhibitor abolished the effect of IL-36 on *PLSCR1* expression (Fig. 6c).

**Figure 6.**
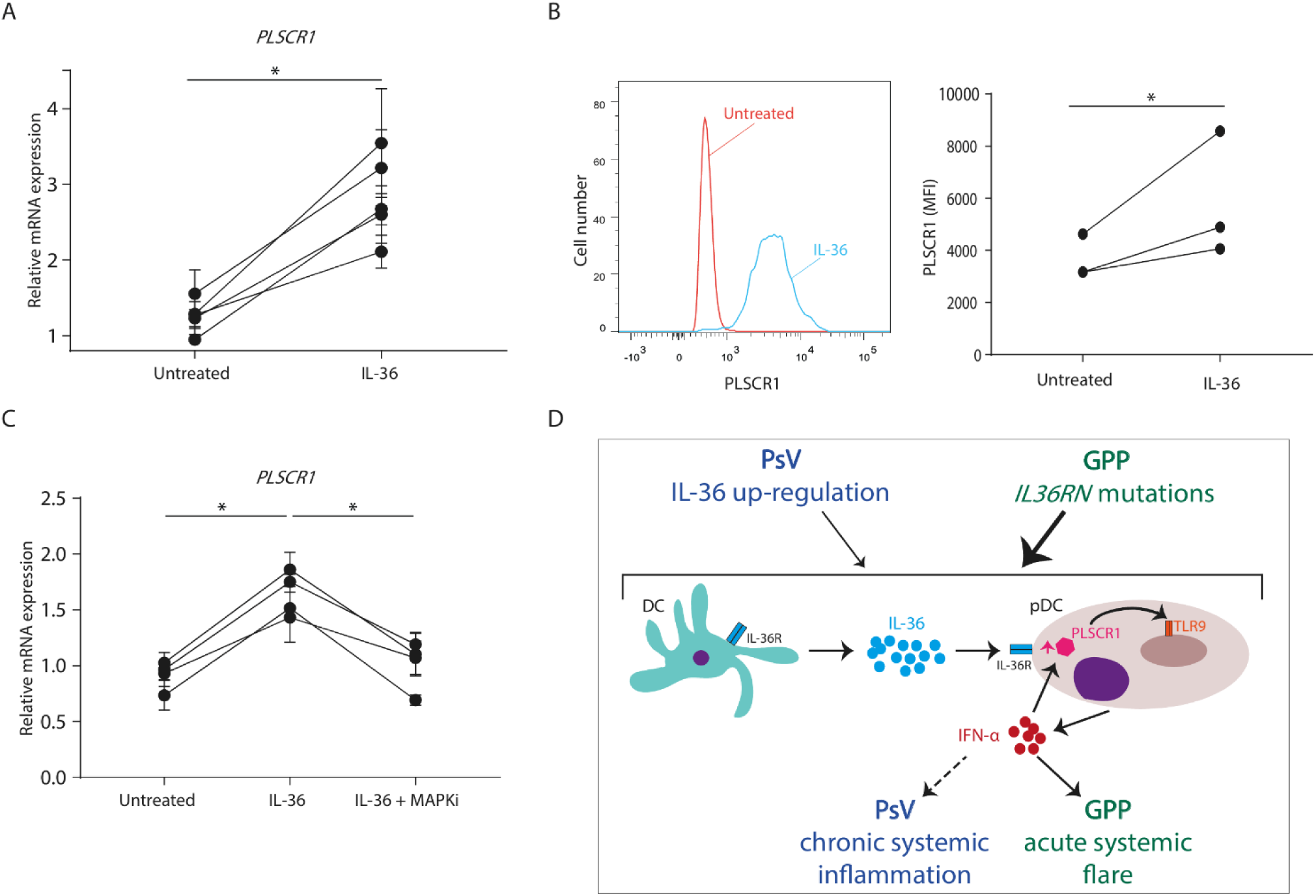
IL-36 up-regulates the expression of PLSCR1. (**A**) Following treatment of PBMCs with IL-36 or vehicle, *PLSCR1* mRNA expression was measured by real-time PCR and normalised to *B2M* levels. Each line in the plot represents an independent healthy donor (n=5). Data points within a line represent mean +/- SEM of results obtained in triplicate stimulations. \**P*<0.05 (Wilcoxon signed-rank test). (**B**) Following IL-36 treatment of PBMCs, PLSCR1 mean fluorescence intensity (MFI) was measured by flow-cytometry, in gated PSLCR1^+^ pDCs. The panel on the left shows a representative histogram, while the plot on the right illustrates the results obtained in individual healthy donors (each is represented by a line). \**P*<0.05 (Wilcoxon signed-rank test). (**C**) Following pre-treatment with SB203580 (MAPKi) or vehicle, PBMCs where stimulated with IL-36. *PLSCR1* expression was then determined by real-time PCR and normalised to *B2M* levels. Each line in the plot represents an independent healthy donor (n=4). Data points within a line represent mean +/- SEM of results obtained in triplicate stimulations. \**P*<0.05 (Friedman’s test with Dunn’s post-test). (**D**) Proposed pathogenic model. Interleukin-36 produced by mDC up-regulates PLSCR1 expression in pDCs, potentiating TLR-9 dependent IFN-α release. IFN-α, in turn, induces further *PLSCR1* transcription, thus propagating an inflammatory feed-forward loop.

Thus, we have demonstrated that IL-36 can act directly on pDCs, where it up-regulates *PLSCR1* transcript and protein levels, in a MAPK-dependent fashion.

## DISCUSSION

The hypothesis underlying this study was that IL-36 signalling de-regulation affects systemic responses in PsV. While the disease has been historically described as a dermatological condition, the importance of extra-cutaneous co-morbidities is increasingly recognised and so is their impact on patient mortality^29^. Of note, the prevalence of most co-morbid disorders increases with the severity and the duration of PsV^9, 30^. There is therefore a dose-dependent association between cutaneous and extra-cutaneous inflammation, which suggests a shared systemic pathogenesis. The underlying pathways, however, remain poorly understood.

Here, we demonstrated that IL-36 signalling is up-regulated in the leukocytes of PsV patients. We also showed that this abnormal activity correlates with a signature of Type-I IFN over-expression. Of note, several studies have found that Type-I IFN plays a key role in vascular inflammation, as it promotes macrophage-endothelial cell adhesion and stimulates the recruitment of leukocytes to atherosclerotic plaques ^31, 32^.

In keeping with these observations, signatures of excessive Type-I IFN activity have been documented in various diseases presenting with prominent systemic involvement. One notable example is systemic lupus erythematosus (SLE), a multi-organ disorder that is associated with accelerated atherosclerosis, skin and joint inflammation^33^. Of interest, three independent studies have reported that IL-36 serum levels correlate with disease activity in SLE^34, 35, 36^, which further reinforces the link between IL-36 and Type-I IFN. Our study adds to these observations and provides mechanistic insights into the underlying inflammatory pathways.

Our computational and experimental results implicate pDCs as the most likely mediators of IL-36 activity. First, the analysis of the GPP RNA-sequencing data identified the activation of IRF7 as one of the most significant drivers of differential gene expression. Second, flow cytometry studies demonstrated that IL-36R surface expression is highest in pDCs, especially among GPP patients. Finally, we showed that IL-36 acts directly on pDCs, where it up-regulates PLSCR1 mRNA and protein expression.

Of note, *PLSCR1* siRNA knockout inhibits type I IFN production by human pDCs^27^, so it is reasonable to hypothesise that an increase in gene expression would have the opposite effect. While the PLSCR1 induction observed in our experiments was modest (IL-36 increased gene expression levels by 1.5-2.0 fold), it might be sufficient to set in motion a feed-forward loop whereby up-regulated PLSCR1 promotes the production of type I IFN, which in turn induces further *PLSCR1* transcription. In fact, self-amplifying loops are a key feature of Type-I IFN signalling, as they are required for the establishment of robust antiviral responses^37^.

We cannot exclude the possibility that additional IL-36 responsive genes or cell types may also contribute to the up-regulation of Type-I IFN. We have found that IL-36 does not affect the expression of *TLR9* or that of key downstream genes (*IRF1, IRF3, IRF7*; data not shown). We have also observed that genes driving other antiviral pathways (*DDX58*/RIG-I, *IFIH1*/MDA5, *TMEM173*/STING) are not systemically up-regulated in PsV or GPP. However, we cannot rule out an effect of IL-36 on as yet undiscovered regulators of innate antiviral immunity.

In conclusion, our computational and experimental studies have identified an IL-36/TLR-9 axis which up-regulates systemic Type-I IFN production in psoriasis (Fig. 6d). In GPP patients, the effects of IL-36 signalling are amplified by inherited *IL36RN* mutations, a phenomenon which is likely to account for the severe nature of systemic flares. In PsV, the Th17-dependent up-regulation of IL-36 cytokines is associated with a less pronounced transcriptional signature and with signs of chronic systemic inflammation. Thus, IL-36 is an important immune regulator in psoriasis, contributing to the systemic pathogenesis of acute and chronic forms of the disease.

Of note, two IL-36 inhibitors are currently in clinical development. Promising results have been reported in GPP Phase I/II trials^38, 39^ and further studies are underway in other conditions thought to be IL-36 mediated (e.g. ulcerative colitis, clinical trial id: NCT03648541). Thus, our study has characterised a novel innate immune axis that could be targeted for the treatment of systemic symptoms in psoriasis.

## METHODS

### Human subjects

The study was performed in accordance with the principles of the Declaration of Helsinki. Patients were ascertained at St John’s Institute of Dermatology and Royal Free Hospital (London, UK), Glasgow Western Infirmary (Glasgow, UK), Salford Royal Foundation Trust (Manchester, UK) and Hospital Sultanah Aminah (Johor Bahru, Malaysia). The study was approved by the ethics committees of participating institutions and written informed consent was obtained from all participants.

Nine unrelated GPP patients (1 male and 8 females, average age: 57) and 7 healthy controls (1 male and 6 females, average age: 52) were ascertained for whole-blood RNA-sequencing, while neutrophil RNA-sequencing was carried out in 8 GPP patients (1 male and 7 females, average age: 51) and 11 healthy controls (1 male and 10 females, average age: 50) (Supplementary Table 1). Five controls were common to both studies, while the overlap in the patient dataset is shown in Supplementary Table 1. A further 35 cases (10 males, 25 females, average age: 49) and 7 controls (6 females and 1 male, average age: 45) were recruited for the validation of whole-blood RNA-sequencing results. For the validation of neutrophil RNA-sequencing, blood was obtained from 17 GPP (2 males and 15 females, average age: 56.5), 26 control (3 males and 23 females, average age: 46.4), 13 APP (3 males and 10 females, average age: 54.3), 9 CAPS (3 males and 6 females, average age: 44.6) and 17 PsV (13 males and 4 females, average age: 45.2) individuals. The key inclusion criterion for the recruitment of patients with PsV was a diagnosis of moderate-to-severe plaque psoriasis (Psoriasis Area Severity Index >10). Individuals receiving anti-TNF treatment, however, were excluded from the study. Blood was also obtained from 4 GPP cases (3 females and 1 male, average age: 49.2) and 4 controls (3 females, and 1 male average age: 33.4) for flow-cytometry assays. Finally, samples donated from 3 healthy volunteers were used for the stimulation of PBMCs and pDCs.

### RNA-sequencing

Total RNA was isolated from whole blood collected in Tempus^TM^ Blood RNA Tube using a Tempus^TM^ Spin RNA Isolation Kit (Thermo). Samples were subjected to globin depletion using a GLOBINclear^TM^ Kit (Thermo). Neutrophil RNA was isolated with GeneJET RNA purification kit (Thermo).

Whole-blood RNA was sequenced on a HiSeq 3000 Illumina platform, obtaining 150bp paired-end reads. Neutrophil RNA was sequenced on a NextSeq 500 Illumina platform obtaining 75bp single-end reads. The quality of the sequence data was assessed using FastQC. Alignment against the HG38 human genome was implemented in TopHat^40^ with indexes generated by Botwie2. Read counts produced by HTseq-count were used as input for the differential expression analysis, which was performed with DESeq2^41^ (R package, v 16.2), using sex as a co-variate.

Genes with a fold change (FC) ≥1.5 and a false discovery rate (FDR) <0.05 were used as input for pathway and upstream regulator enrichment analyses (IPA, Qiagen). The latter assesses the over-representation of targets of known transcription regulators, while also building gene-target networks based on published co-expression and binding affinity data. Here, STAT1-STAT3-and IRF7-centered networks were visualised with the igraph v1.0.1 R Package.

The python library published by Li et al^17^ was modified in order to identify blood transcriptional modules that are enriched among genes up-regulated in GPP. Briefly, the modules that were active in our datasets were selected using the genetable_to_activityscores function. Next, the enrichment_test function was applied to the lists of up-regulated genes, taking into account the module activity scores and fold change of the genes mapping to each module. Enrichment P values were then calculated and corrected for multiple testing using the Benjamini-Hochberg method.

The interferon score was built using the five Type-I IFN dependent genes that were most up-regulated in GPP whole-blood (*PLSCR1, OALS, IFI6, IFIT3, IFITM3)*. As IL-36 dependent genes have not been systematically characterised in leukocytes, the IL-36 score was based on the analysis of five genes which were strongly induced by IL-36 in keratinocytes^3^ and robustly expressed in whole-blood (*IL1B, PI3, VNN2, TNFAIP6, SERPINB1* and *PLSCR1)*. Both scores were derived by normalising RPKM values to a calibrator sample and then computing the median expression of the five signature genes.

### Cell isolation and culture

Neutrophils were purified using the MACSxpress Whole Blood Neutrophil Isolation Kit (Miltenyi Biotec). PBMCs were isolated using Ficoll-Paque PLUS (GE Healthcare). Plasmacytoid dendritic cells were purified from PBMCs using a Plasmacytoid Dendritic Cell Isolation Kit (Miltenyi Biotec). PBMCs and pDCs were cultured at a density of 2.5×10^6^ cells/ml and 2.5×10^5^ cells/ml, respectively, in RPMI Glutamax (Gibco) supplemented with 10% FBS and 1% penicillin-streptomycin. Cells were stimulated with 50 ng/ml IL-36α (Bio-Techne) for 6 hours and with 1.6 ng/ml ODN-A CpG (Invivogen) for a further 6 hours. For IFN-α and PLSCR1 flow cytometry analysis, Brefeldin A (BioLegend) was added to the stimulated cells at a 1: 1,000 dilution after 9 hours. Response to stimulation was measured by real-time PCR, ELISA or flow-cytometry.

### Real-time PCR and ELISA

RNA samples were isolated with the GeneJET RNA purification kit (Thermo). Following reverse transcription with the nanoScript2 kit (Primerdesign), gene expression was assessed by real-time PCR using PrecisionPLUS Master Mix with SYBR and ROX (Primerdesign) in conjunction with the primers listed in Supplementary Table 2. The IFN score was derived by computing the median RQ of the five signature genes, (*PLSCR1, OALS, IFI6, IFIT3, IFITM3)* according to the method described by Rice et al^42^. The production of IFN-α was measured using the Human IFN-alpha ELISA kit (Bio-Techne). For whole blood and PBMC samples, transcript levels were normalised to *B2M* expression, while *RPL13A* was used for neutrophils.

### Flow cytometry

The purity of neutrophil isolated for RNA-sequencing was measured by staining cells with anti-CD45, anti-CD15, anti-CD16, anti-CD3, anti-CD24 and anti-CD19 antibodies. IL36R surface expression and IFN-α levels were measured by staining PBMCs with LIVE/DEAD^TM^ Fixable Near-IR (Invitrogen), Fc and monocyte blocker (Biolegend), antibody against the protein of interest and an antibody cocktail for monocytes (anti-CD3, anti-CD20, anti-CD19, anti-CD16, anti-CD14, anti-CD56) or DCs and ILCs (Lineage, HLA-DR, CD123, CD11c, CD127). IL36R expression on neutrophils was determined by staining for the receptor as well as CD15, CD16 and CD14. PLSCR1 expression was measured by staining purified pDCs with anti-CD123, anti-HLA-DR, anti-CD11c and anti-PLSCR1. Cells were acquired on a BD Fortessa LSR or a BD FACSCanto II instrument. All data was analysed using FlowJo v10 software. The details of all antibodies are reported in Supplementary Table 3.

### Statistics

The tests used in each experiment are reported in the legend of the relevant figure. Briefly, differences between the cytokine scores of patient and control groups were assessed using a two-tailed t-test or one-way ANOVA, as appropriate. To account for donor variability in cytokine responses, the results of IL-36/CpG stimulations were analysed with non-parametric methods, as these do not assume equal variance among samples. Thus, Wilcoxon signed-rank test was used for comparisons between two groups and Friedman’s test for comparisons between three groups. The correlation between cytokine scores was calculated using the Spearman method. The significance of the overlaps observed in Venn diagrams was computed with a hyper-geometric test. A two-tailed Fisher’s exact test was used to compare the clinical features of patients with high and low IFN scores. Unless otherwise indicated, experimental data are shown as means +/-SD and p-values < 0.05 were considered as statistically significant.

## DATA AVAILABILITY

The RNA-sequencing datasets generated during the study have been submitted to the Gene Expression Omnibus (GEO). The unique identifier is GSE123787. Supplementary data will be available on request.

## ACKNOWLEDGEMENTS

We are very grateful to Dr Paola Di Meglio for her helpful comments and technical advice. We also wish to thank the patients and volunteers who took part in this study. We acknowledge support from the Department of Health via the National Institute for Health Research (NIHR) BioResource Clinical Research Facility and comprehensive Biomedical Research Centre award to Guy’s and St Thomas’ NHS Foundation Trust in partnership with King’s College London and King’s College Hospital NHS Foundation Trust (guysbrc-2012-1). The APRICOT clinical trial is funded by the Efficacy and Mechanism Evaluation (EME) Programme, an MRC and NIHR partnership (grant EME 13/50/17 to CHS, FC and JNB). This study makes use of data generated by the Blueprint Consortium. A full list of the investigators who contributed to the generation of the data is available from www.blueprint-epigenome.eu. Funding for the project was provided by the European Union’s Seventh Framework Programme (FP7/2007-2013) under grant agreement no 282510 – BLUEPRINT. MC is supported by the Psoriasis Association, MV by the UK Medical Research Council and SKM by a NIHR Clinical Lectureship.

The views expressed in this publication are those of the author(s) and not necessarily those of the MRC, NHS, NIHR or the Department of Health.

## AUTHOR CONTRIBUTIONS

FCa designed the study and supervised the experimental work. MC, MV and MR carried out the experiments, with input from GL for the flow cytometry analysis. MC, MV, IMC and FCi implemented the RNAseq and the downstream data analysis. SKM, ADB, SEC, HSY, CHS and JNB coordinated the ascertainment and clinical characterisation of patients, as well as the recruitment of healthy volunteers. FCa wrote the manuscript with input from GL, SKM, CHS, FCi and JNB.

